# PubMind: Literature-Based Genetic Variant Extraction and Functional Annotation Using Large Language Models

**DOI:** 10.1101/2025.10.13.682183

**Authors:** Peng Wang, Kai Wang

**Affiliations:** Center for Cellular and Molecular Therapeutics, Children’s Hospital of Philadelphia, Philadelphia, PA 19104, USA; Department of Pathology and Laboratory Medicine, University of Pennsylvania, Philadelphia, PA 19104, USA

**Author notes:** Correspondence should be addressed to K.W.

## Abstract

The rapid growth of biomedical literature has produced extensive functional knowledge on genetic variants, much of which remains buried in unstructured texts. Current databases such as ClinVar and the Human Gene Mutation Database (HGMD) attempt to catalog this knowledge but have significant limitations: ClinVar depends on voluntary submissions and covers only a fraction of published literature, while the academic version of HGMD is updated infrequently and provides limited functional annotation. To address these gaps, we developed PubMind, an AI-driven multi-layer framework that uses large language models (LLMs) to extract variant– function–disease associations and supporting evidence from text. PubMind integrates a fine-tuned BERT model for input triage with instruction-tuned GPT models for inferring disease associations and functional annotations. The system captures diverse variant types—including SNVs, CNVs, SVs, and gene fusions—and normalizes records to genome and transcriptome coordinates. Benchmarking demonstrates >90% accuracy in variant recognition and 99% precision in disease extraction. Application of PubMind on >41 million PubMed abstracts and >5 million open-access full-text articles produced PubMind-DB, a database containing ∼1.3 million unique variants with rich contextual annotations, accessible via a web interface and API. Only ∼10% of PubMind’s variants overlapped with ClinVar entries, yet >80% showed concordant pathogenicity labels, including full agreement with ClinVar’s expert-reviewed variants. Case studies demonstrate PubMind-DB’s ability to uncover supporting evidence for variant pathogenicity that might otherwise be missed by manual searches. Together, these findings establish PubMind as a scalable LLM-based framework that transforms unstructured biomedical text into structured genomic knowledge, advancing variant interpretation for precision medicine.

## Introduction

The exponential growth of biomedical research has yielded a vast corpus of knowledge on human genetic variants and their disease associations. Yet much of this information remains locked in unstructured text, inaccessible to automated systems to catalog such variants. Existing resources, such as ClinVar ^1^ and Human Gene Mutation Database (HGMD) ^2^, attempt to catalog variants but they both face critical limitations. ClinVar depends on voluntary submissions, leading to heterogeneity in quality, limited coverage of published literature, and frequent absence of disease or phenotype annotations—even for variants labeled Pathogenic or Likely Pathogenic (P/LP). Nevertheless, ClinVar provides great value to the understanding of genetic variants and has been routinely used as benchmark dataset. Additionally, some of its limitations, such as the lack of confidence and phenotype annotations, are gradually being addressed by community efforts such as ClinVar STAR system and ClinGen consortium ^3^, even though such efforts are still labor intensive with limited coverage of genome. On the other hand, the HGMD directly uses literature as input and provides variant-disease association, through manual review and maintenance. However, the academic (publicly available) version of HGMD is infrequently updated, lags behind the professional subscription version by years (for example, the last public version update in December 2021 vs. professional version update in April 2024), and covers only ∼60% of curated entries ^2^. Moreover, HGMD records provide only variant–disease association with a reference, yet in many cases it does not specify the nature of such association (e.g., benign vs. pathogenic), and it does not provide functional/experimental evidence which is necessary to assist in variant interpretation ^4^.

A significant portion of variant-related knowledge, including subtle pathogenicity assertions and mechanistic insights, remains buried within full-text articles that are rarely curated into structured databases. To overcome the challenge of variant extraction from text, conventional biomedical Named Entity Recognition (NER) systems, such as tmVar3.0 ^5^, PubTator3 ^6^, AIONER ^7^, GNormPlus ^8^, and GNorm2 ^9^, have improved gene, variant and disease tagging. But they are largely limited to entity-level extraction and lack the capacity to recover relational and context-dependent information, such as variant-specific pathogenicity in different disease contexts or experimental outcomes supporting these claims. Rule-based heuristics further limit adaptability, requiring constant manual updates to keep pace with evolving nomenclature.

Further effort has been made through information aggregation system, such as LitVar ^10^, to link literature paragraph with variant information from different sources. LitVar is a web interface which connects variant records found in literature paragraph, extracted by tmVar3.0 ^5^, with external variant database for populational information and pathogenicity such as dbSNP ^11^. LitVar greatly accelerates the variant knowledge retrieval through providing a web interface that connects difference sources of information together, trying to enrich the limited functional and pathogenic information extracted by the conventional NER systems. However, the variant information in LitVar is still limited by the extraction efficiency and accuracy of external extraction tool, and relies on the existing variant record in external datasets. LitVar does not extract variant–disease-pathogenicity associations directly from text, nor do they resolve contextual semantics or experimental evidence. For example, searching for BRCA1 p.Cys61Gly (p.C61G) mutation in LitVar2 only provided pathogenicity and allele frequency info based on the matched dbSNP record ^11^ alongside raw PubMed text snippets. To understand which paper considers this variant as pathogenic, one needs to read the paragraph from every PubMed article. Therefore, this procedure may not be optimal for quickly summarizing the mutational landscape of a gene in a specific disease.

The advent of a new generation of AI-based methods for unstructured data, such as Large Language Models (LLMs) ^12–14^, presents a transformative shift in this landscape. Advances in model architecture and input token capacities, exemplified by the open-source LLaMA 3 family ^15^, have enabled LLMs to process full-text articles with nuanced contextual understanding. Unlike conventional NER pipelines that focus on isolated entities, LLMs are capable of context-aware reasoning, leveraging few-shot learning to infer complex relationships among variants, diseases, and phenotypes—even when such associations are implied rather than explicitly stated. For example, the missense variant p.F252L in *IRF6* was predicted by multiple computational tools to be benign, yet functional assays in maternal-null irf6–/– zebrafish revealed its inability to rescue the rupture phenotype ^16^. Sophisticated LLMs can correctly infer this variant as “likely pathogenic”, prioritizing experimental evidence over misleading computational predictions. This represents a paradigm shift from syntactic extraction toward context-aware comprehension of variant knowledge.

Recent advances with domain-adapted transformer models such as BioBERT ^17^, PubMedBERT ^18^, PhenoBCBERT ^19^, and ClinicalBERT ^20^ further underscore the potential of LLMs in biomedical text mining. These models outperform traditional tools on relation extraction tasks, particularly when fine-tuned on curated corpora or refined with human-in-the-loop strategies. However, BERT-based systems remain constrained by their classification architecture: each task requires separate fine-tuning, limiting generalizability and scalability. They also lack native support for multi-turn instruction-following and show limited compositional reasoning beyond their training domain.

To address these limitations, we developed **PubMind**: **Pub**lication **M**utation and **in**formation **d**iscovery using LLMs. PubMind represents a novel, scalable pipeline that harnesses the generative and reasoning capabilities of GPT-style models, specifically leveraging the LLaMA 3.3 architecture, for end-to-end variant extraction from both abstracts and full-text articles. Our workflow integrates a fine-tuned BERT triage stage with downstream GPT-based inference and normalization modules for variant annotation, genome coordinate resolution, and disease/phenotype mapping to standardized ontologies such as MONDO ^21^ and HPO ^22^. PubMind provided enriched functional and pathogenic information of variants, which is extracted directly from literature with evidence provided by LLM reasoning. PubMind was able to provide literature-specific annotation that LitVar2 did not provide: Searching the same BRCA1 p.Cys61Gly (p.C61G) mutation, we found 79 records across multiple publications for the same variant (PubMind ID: PVID117180), so that we can consolidate pathogenicity assignments, maps diseases to MONDO terms ^21^ to standardize disease name description, and provides human interpretable reasoning (e.g., “abolishes BRCA1 interaction with BARD1”).

Through application on 41.7 million PubMed abstracts and 5.4M full texts, we further developed PubMind-DB, a database containing ∼1.3 million unique variants with rich contextual annotations, accessible via a web interface and API. By supporting complex variant types (SNVs, gene fusions, SVs, and CNVs) and offering a continuously updated, user-accessible knowledgebase, PubMind bridges the gap between unstructured biomedical text and structured genomic interpretation, advancing variant interpretation and accelerating applications in precision medicine. We note that PubMind can be applied to institutionally licensed literature resources to build private versions of variant databases by users. Finally, PubMind empowers healthcare systems to construct secure, institution-specific variant interpretation databases directly from clinical notes and reports, thereby facilitating the implementation of genomic medicine on scale.

## Results

### An overview of the PubMind workflow

We developed PubMind, a large language model (LLM)-assisted framework for *Publication Mutation and information discovery*, designed to extract variant–disease–pathogenicity relationships directly from biomedical literature. We also used PubMind to process 41,682,357 PubMed abstracts and 5,425,084 full-text articles from PMC open-access subset, and transformed unstructured text into a structured, searchable knowledgebase called PubMind-DB (**Figure 1a**). At its core, PubMind integrates multiple components into a scalable, multi-layer pipeline optimized for variant interpretation.

**Figure 1.**
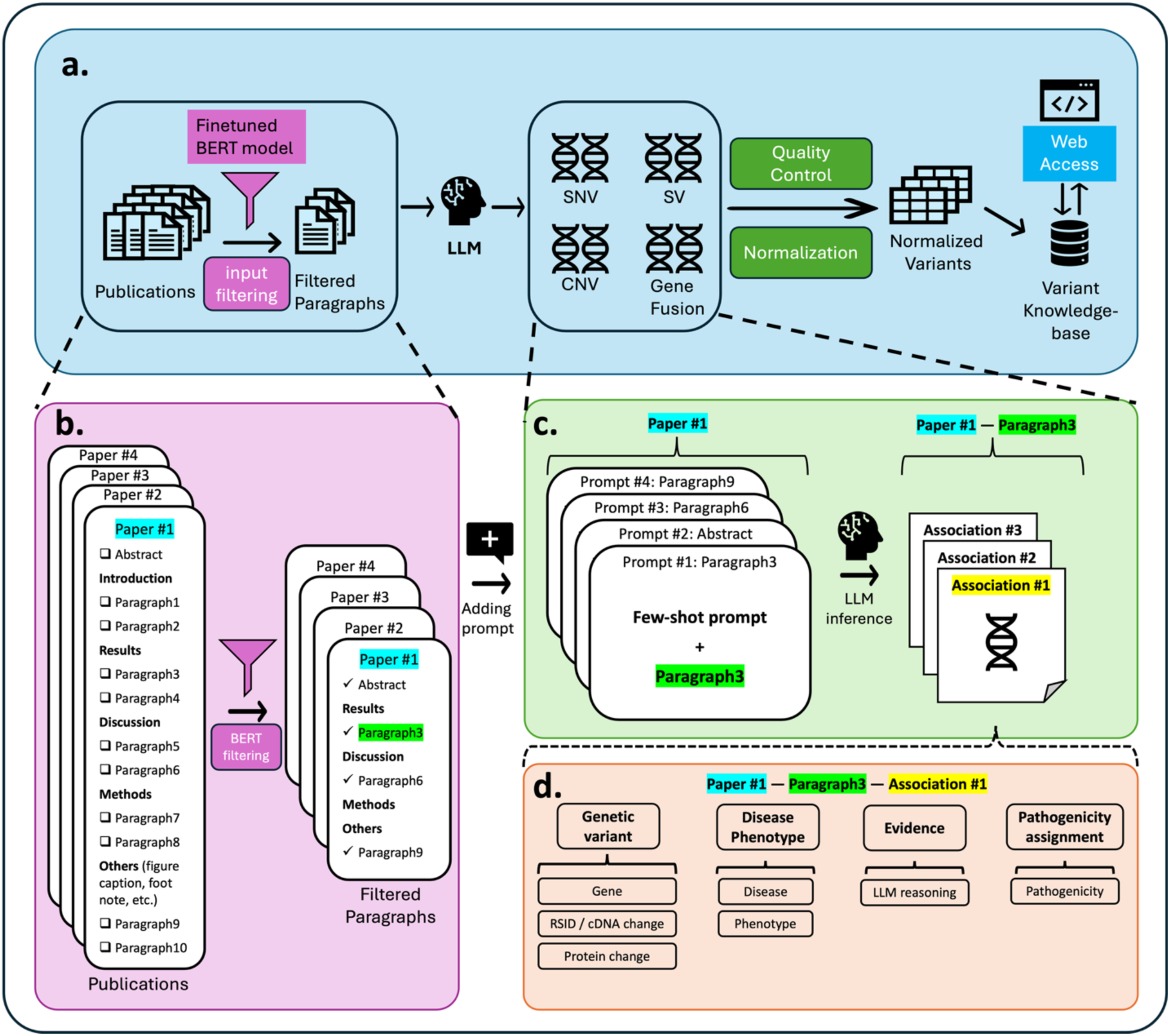
Overview of the PubMind architecture. (a) PubMind integrates a multi-stage workflow for large-scale extraction of variant–disease–pathogenicity associations. Publications are first processed through a fine-tuned BERT filtering module, which reduces input size by retaining only paragraphs enriched for genetic variant information. Filtered text is then passed to an instruction-tuned LLM for extraction. The inferred variant records are subsequently validated, normalized, and stored in the PubMind database (PubMind-DB), which is accessible through a web interface for querying, visualization, and data export. (b) Filtering module. PubMed abstracts and PMC full-text articles are segmented by section, and a fine-tuned BERT model selects paragraphs containing relevant gene, variant, disease, or pathogenicity content. (c) Inference module. Retained paragraphs are processed with structured prompts tailored to different variant types, enabling the LLM to extract standardized attributes (gene, variant representation, disease, phenotype, pathogenicity, and reasoning). (d) Example association. An illustration of how a paragraph yields a structured variant–disease–pathogenicity association, including variant identifiers (RSID, cDNA, protein change), mapped disease/phenotype terms, supporting textual evidence, and pathogenicity assignment.

The workflow begins with a fine-tuned BERT model that filters and prioritizes abstracts or paragraphs enriched for mentions of genes, variants, diseases, and pathogenicity (**Figure 1b**). These candidate texts are then passed through a set of carefully engineered prompts, each tailored to extract different attributes for specific variant classes—single nucleotide variants (SNVs), copy number variants (CNVs), structural variants (SVs), and gene fusions. For each input, a large instruction-tuned LLM (LLaMA3.3-70B) infers the semantic relationships and extracts variant–disease–pathogenicity associations directly from the text, uncovering biological meaning that is often buried (**Figure 1c–d**). Unlike traditional entity recognition methods, PubMind does not rely on manual rule sets or model retraining, allowing it to generalize across diverse literature. Extracted results are post-processed, normalized by position (coordinate) and disease/phenotype, and stored in a relational database, where they undergo validation procedures (see **Methods** and **Supplementary Methods**). The curated database is then exposed through a web-based interface that supports flexible querying, visualization, and export of results, enabling users to rapidly access literature-derived variant insights. This modular architecture makes PubMind interpretable, scalable, and adaptable to new biomedical domains, providing a foundation for comprehensive literature-based variant annotation.

### Benchmarking and assessment of PubMind

To rigorously assess PubMind’s performance, we first focused on the SNV extraction task as a representative benchmark. We evaluated several LLaMA and DeepSeek-R1-series models for accuracy in variant recognition (**Supplementary Fig. 1**). Considering the computational efficiency and memory resource requirement, LLaMA3.3-70B demonstrated the highest precision for both cDNA- and protein-level variants and was selected as the primary inference model. Next, we compared prompting strategies—including zero-shot, chain-of-thought (CoT), and few-shot prompting (**Supplementary Table 1**). A short system prompt with few-shot learning achieved the best balance, yielding >90% accuracy in SNV extraction while maintaining efficient GPU runtime (**Supplementary Fig. 2a–b**). Using the same prompt, PubMind also extracted disease names with 99.9% accuracy without hallucinations (**Supplementary Fig. 3c**).

Pathogenicity assignment in PubMind is inferred directly from literature context, and may not fully align with ClinVar classifications. Thus, we performed benchmarking against ClinVar to assess concordance and examine reasons when differences arise. Across all PubMind-DB SNVs, there are 1,017,540 literature-extracted variant records which are consolidated into 579,844 unique variants. From the consolidated unique variants, 419,012 unique variants were mapped to transcripts; after accounting for multiple codon-level representations, this resulted in 916,538 transcript-mapped genomic variants (VCF format), which were annotated using ANNOVAR ^23^. Among these, 97,214 genomic variants (10.6%) overlapped with ClinVar records (**Fig. 2a**). The relatively small overlap highlighted the advantage of PubMind in cataloging literature-derived variants, while the observed concordance patterns illustrated its reliability.

**Figure 2.**
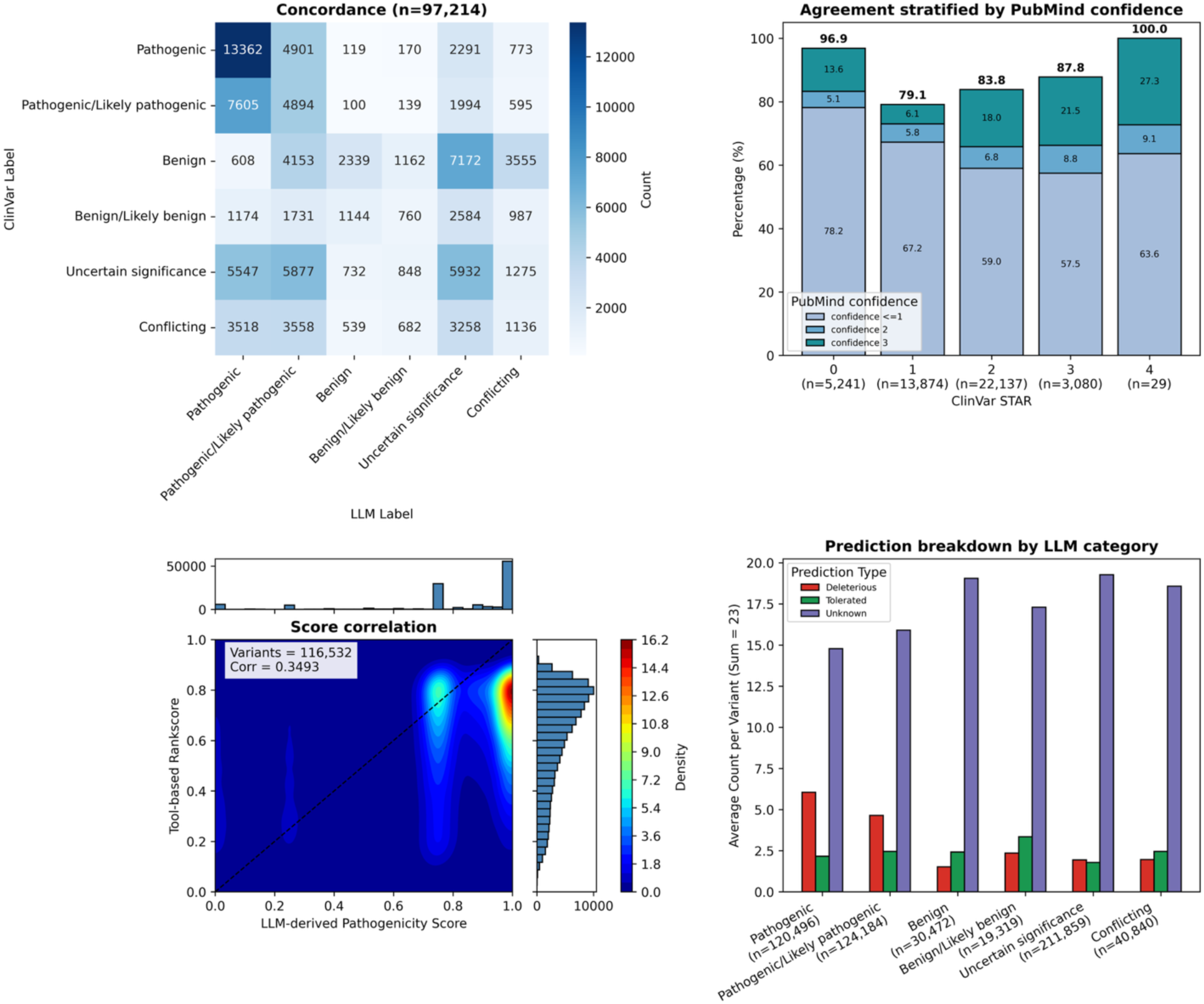
Benchmarking PubMind pathogenicity assignments against ClinVar and bioinformatics tools. (a) Concordance with ClinVar. Heatmap showing the overlap between PubMind-derived pathogenicity classifications and ClinVar labels (n=97,214). Agreement is highest for pathogenic variants, while benign annotations show greater heterogeneity. (b) Stratification by ClinVar review status. Agreement between PubMind and ClinVar increases with review depth, reaching 100% concordance for four-star variants. Stacked bars indicate PubMind confidence scores, which also rise with higher ClinVar review status. (c) Score correlation. Comparison of PubMind-derived pathogenicity scores with the average rankscore from 51 computational tools. Variant density is represented by a continuous heatmap, with warmer colors indicating regions of higher concentration. Marginal histograms show the distribution of scores for PubMind (top) and computational tools (right). Variants predominantly cluster at high pathogenicity values, though PubMind additionally identifies low-scoring benign variants that are less represented by computational tools. (d) Tool-based categorical predictions. Average predictions from 23 categorical bioinformatics tools across different PubMind pathogenicity categories. Variants classified as pathogenic/likely pathogenic by PubMind align with higher deleterious predictions, whereas benign categories align with fewer pathogenic calls. Most tool predictions fall into the “unknown” category.

Agreement with ClinVar was the highest for pathogenic variants, but lower for benign classifications—consistent with known variability in annotating benign variants between different tools. To refine this comparison, we stratified concordance by ClinVar expert review status (“Star” system), which reflects confidence in classification (**Fig. 2b**). Agreement between PubMind and ClinVar increased with review depth, reaching 100% concordance for four-star variants. Importantly, PubMind confidence scores also tracked with ClinVar Stars, indicating that variants curated more thoroughly in ClinVar were also classified with higher confidence by PubMind. This trend underscored the robustness of PubMind’s pathogenicity assignments, while suggesting that ClinVar 0-star entries—often heterogeneous in quality—account for most discrepancies.

We next compared LLM-derived pathogenicity scores with consensus predictions from diverse bioinformatics tools using density plot (**Fig. 2c**). This tool-based score is the average pathogenicity score (rank from 0 to 1) from 51 bioinformatics tools prediction using dbNSFP v4.7 ^24^. PubMind-derived scores ranged from 0 (benign) to 1 (pathogenic) with peaks around 0.75 and 1.0, while tool-based scores displayed a distribution skewed toward ∼0.8. Both methods showed strong agreement in the high-pathogenicity range, with dense clustering in the top-right quadrant of the plot. However, PubMind additionally captured low-scoring peaks corresponding to benign calls in literature, whereas bioinformatics tools tended to yield heterogeneous or high scores. This suggests that PubMind provides contextual evidence not consistently captured by computational predictors. Finally, we compared categorical predictions from 23 bioinformatics tools in dbNSFP against PubMind classifications (**Fig. 2d**). As expected, variants labeled as pathogenic/likely pathogenic by PubMind showed a higher proportion of deleterious predictions, while benign/likely benign variants aligned with lower proportion of deleterious calls by bioinformatics tools. However, the majority of output from bioinformatics tools were “unknown,” reflecting the absence of confident classifications. PubMind, by contrast, was able to provide structured pathogenicity insights directly from text, complementing predictions by bioinformatics tools and offering annotations for variants otherwise left unclassified.

### Exploring the PubMind-DB knowledgebase

PubMind-DB encompasses four major variant classes—single nucleotide variants (SNVs), gene fusions, structural variants (SVs), and copy number variants (CNVs)—with standardized identifiers (PVID) and literature-derived annotations (**Table 1**). SNVs are indexed using gene names with corresponding cDNA, RSID, or amino acid changes, yielding 579,844 consolidated unique SNVs, of which 419,012 have genomic coordinates. Gene fusions are defined by their fusion partners and affected protein domains, consolidating into 85,966 fusions from 174,784 literature-derived records. SVs and CNVs are indexed by gene names, chromosomal regions, and coordinates where available, with literature-based disease and pathogenicity associations extracted for each. In total, PubMind catalogs 69,993 unique SVs consolidated from 191,643 literature-extracted SV records, and 29,457 unique CNVs consolidated from 89,953 literature-extracted CNV records, creating a unified resource across diverse variant types.

**Table 1.** Overview of variant records in PubMind-DB. Summary of variant classes extracted from PubMed abstracts and full-texts, after multi-stage processing (literature-derived records, consolidated unique variants, transcript/genomic mapping). ClinVar overlap is reported for transcript-mapped SNVs, based on ClinVar version number 20240914.

| Variant type | Literature-derived records | Consolidated unique variants (unique PVIDs) | Genomic/transcript-mapped variants (VCF-format) | ClinVar overlap (n, %) |
| --- | --- | --- | --- | --- |
| SNV | 1,017,540 | 579,844 | 916,538 | 97,214 (10.6%) |
| Gene fusion | 174,784 | 85,966 | – | – |
| Structural variant (SV) | 191,643 | 69,993 | – | – |
| Copy number variant (CNV) | 89,953 | 29,457 | – | – |
| **Total** | **1,473,920** | **765,260** | **916,538** | **97,214 (10.6%)** |

Figure 3 provides an overview of the pathogenicity label, confidence score, and disease associations represented in PubMind-DB. As shown in Figure 3a, pathogenic and likely pathogenic variants dominate across all variant types, with relatively fewer benign or likely benign annotations. Each record is further assigned a confidence score reflecting the depth of supporting evidence extracted from literature (**Supplementary Table 2**). As expected, the number of variants decreases with increasing confidence level (Fig. 3b). To explore disease context, we identified the top 10 most frequently mentioned diseases for each variant class, stratified by pathogenicity (Fig. 3c). These distributions reveal that most variants are annotated as pathogenic or of uncertain significance (“Other”), while benign classifications are comparatively rare in the published records.

**Figure 3.**
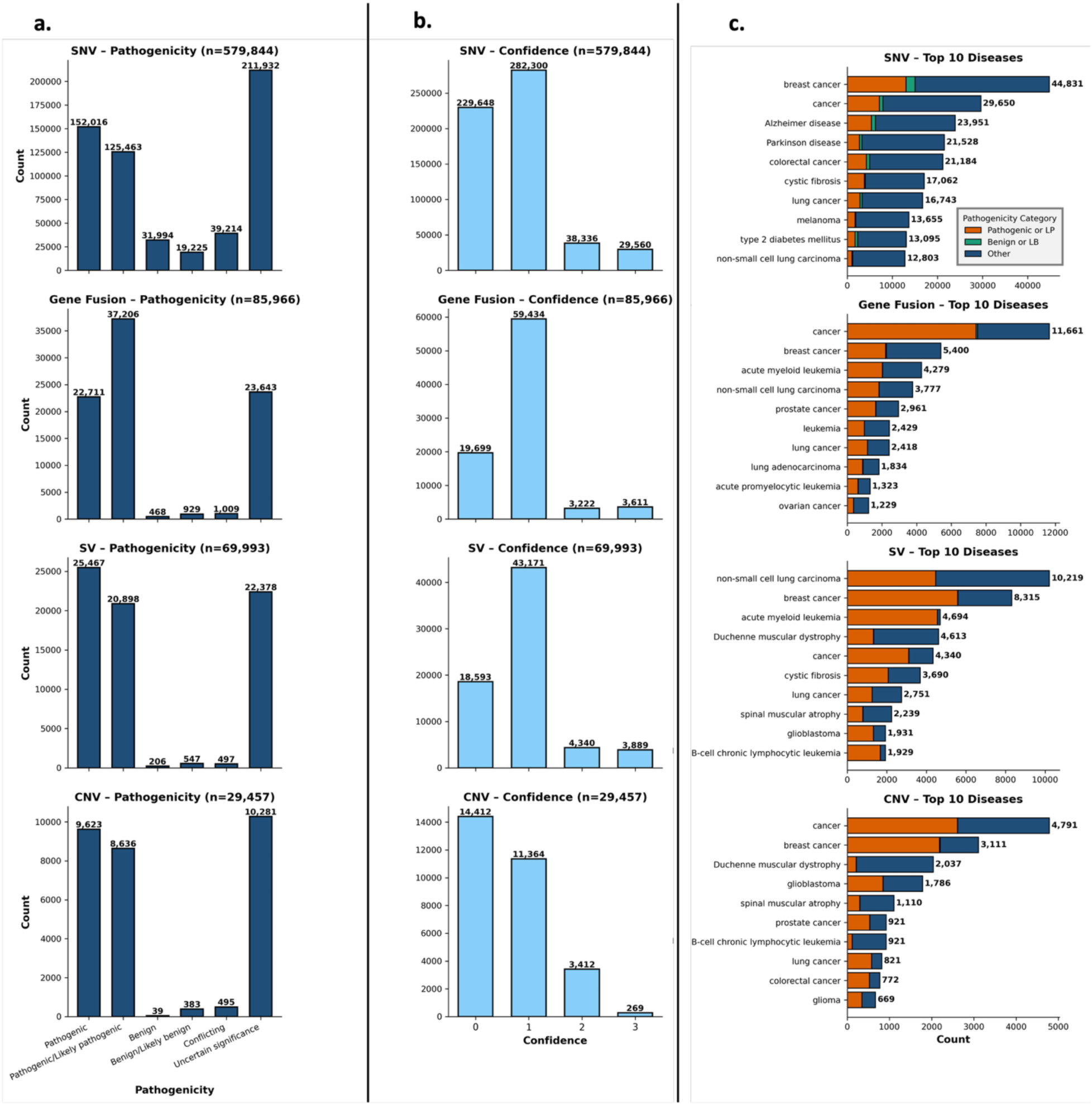
Pathogenicity distribution, PubMind confidence scores, and disease associations across variant types. (a) Pathogenicity classification. Distribution of PubMind-assigned pathogenicity labels for single nucleotide variants (SNVs), gene fusions, structural variants (SVs), and copy number variants (CNVs). Pathogenic and likely pathogenic annotations dominate across all variant types, whereas benign classifications are comparatively rare. (b) Confidence scores. Variants stratified by PubMind confidence score (0–3), reflecting evidence depth and annotation reliability. Most variants are supported at lower confidence levels, with progressively fewer variants at higher tiers. (c) Top 10 disease associations. The ten most frequently mentioned diseases for each variant type, with bars colored by PubMind-assigned pathogenicity category (Pathogenic/Likely Pathogenic, Benign/Likely Benign, or Other [unknown/conflicting]). The majority of top disease associations are driven by pathogenic or uncertain classifications.

We further examined the literature sources underlying PubMind annotations (**Supplementary Fig. 4**). Most variants were extracted from sections classified as “Other” in the XML structure, reflecting text outside standard categories such as Abstract, Introduction, or Result (**Suppl. Fig. 4a**). Across variant types, the temporal distribution of publications peaked around 2021, consistent with the rapid growth of sequencing and genomics literature in recent years (**Suppl. Fig. 4b**). Finally, we identified the top contributing journals for each variant class, with PLoS One and Scientific Reports consistently ranking among the largest sources of variant records (**Suppl. Fig. 4c**).

Together, these results illustrate the breadth and richness of the PubMind database, spanning millions of literature-mined variant annotations with structured pathogenicity and disease associations. By integrating multiple variant classes, a confidence scoring framework, and contextual annotations, PubMind provides a comprehensive, literature-grounded complement to human curated databases, enabling researchers to explore variant knowledge at an unprecedented scale.

### Case Studies: From Variant Reclassification to Therapeutic Context with PubMind-DB

To illustrate use cases of literature-driven pathogenicity annotation, we performed a case study on variants of uncertain significance (VUS) in the *IRF6* gene, a key locus implicated in orofacial cleft. In prior work, Edward et. al experimentally evaluated 37 IRF6 missense mutations using a phenotype rescue assay in irf6−/− zebrafish ^16^. Here, we focused on 9 missense mutations reported in full-text literature ^16^ and compared their pathogenicity assignments across multiple sources: ClinVar ^1^, HGMD ^2^, AlphaMissense ^25^, an ensemble of 21 bioinformatics tools ^26^, and PubMind (Fig. 4).

**Figure 4.**
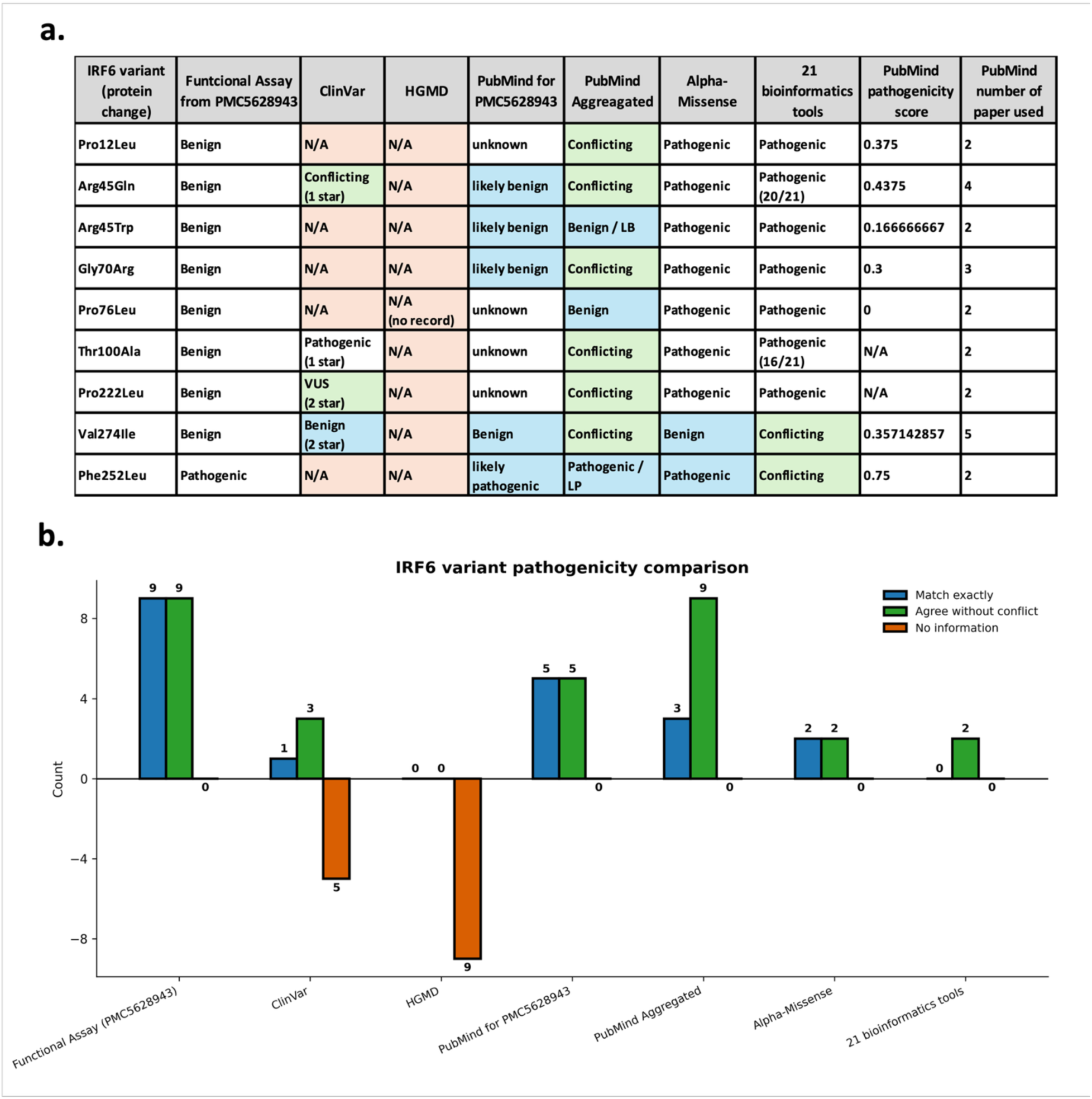
Case study of IRF6 VUS reclassification using PubMind. (a) Comparison of pathogenicity assignments. Pathogenicity calls for nine IRF6 missense variants are shown across multiple sources, including ClinVar, HGMD, AlphaMissense, an ensemble of 21 bioinformatics tools, and PubMind (single-paper and aggregated annotations). Results are benchmarked against functional rescue assays in irf6 −/− zebrafish. Background colors indicate: blue = exact match with functional assay result; green = agreement without conflict (exlucding exact match); orange = no available information (N/A). (b) Summary of concordance. Visualization of the comparison in panel (a). Bars show the number of variants with exact matches (blue), agreements without conflict (green), or missing information (orange). PubMind aggregated annotations achieved complete concordance (9/9), outperforming ClinVar, HGMD, AlphaMissense, and bioinformatics tool predictions.

HGMD provided only variant–disease associations without explicit pathogenicity calls; thus, pathogenicity was unavailable (N/A) for these entries, even though 8 of the 9 variants were indexed. For PubMind, we examined both the annotations derived from this exact zebrafish study (PMCID: PMC5628943 ^16^) and the aggregated annotations across multiple publications. Among the nine variants, eight were functionally rescued *in vivo*, indicating a benign classification. PubMind was the only resource that achieved complete concordance (9/9) when aggregating evidence across publications, correctly identifying all rescued variants as benign or non-pathogenic. Notably, PubMind’s aggregated annotations resolved four variants that remained “unknown” when relying on the zebrafish study alone. In contrast, AlphaMissense ^25^ correctly classified only one of the eight benign variants, and the 21-tool ensemble approach failed to classify any as benign.

Our second use case focused on *PDGFRB*, a receptor tyrosine kinase whose activating variants are well-established drivers of infantile myofibromatosis and other disorders, and for which targeted inhibitors such as imatinib are clinically effective ^27, 28^. In a prior collaborative study, we attempted to search published case studies on *PDGFRB* variants with treatment by imatinib, and manually identified seven *PDGFRB* variants through this exercise. Using PubMind-DB, we systematically queried all reported PDGFRB variants and cross-referenced the “LLM reasoning” column for therapeutic context. All four SNVs previously identified were recovered by PubMind-DB, along with additional clinically relevant variants such as Arg370Cys (PVID702484) and Asn666Ser (PVID702494). Importantly, for two recurrent SNVs (PVID702492 and PVID702578), the LLM reasoning explicitly captured therapeutic evidence, e.g., “clinical improvement when treated with imatinib, an inhibitor of several kinases” and “response to imatinib treatment”. This highlights PubMind-DB’s ability to extract not only variant–disease–pathogenicity associations, but also actionable treatment insights directly from literature.

Beyond SNVs, PubMind-DB proved especially powerful in finding gene fusions and SVs with therapeutic relevance. Across 339 *PDGFRB*-associated fusions found by PubMind-DB, 45 contained reasoning that directly mentioned imatinib. Examples include PDGFRB::EBF1, PDGFRB::ETV6, and COL1A1::PDGFRB, each linked by LLM reasoning to imatinib responsiveness. For instance, the entry for COL1A1::PDGFRB (PVID-GF17305) noted: “The presence of the COL1A1–PDGFRB fusion leads to activation of the PDGFRB tyrosine kinase, which can be targeted by specific inhibitors such as imatinib.” Similarly, 5 of 31 SVs in *PDGFRB* were annotated with imatinib in their reasoning fields. Together, these findings demonstrate how PubMind-DB can consolidate dispersed evidence into a single knowledgebase, enabling rapid retrieval of both pathogenic variants and associated therapeutic opportunities.

## Discussion

In this work, we address the challenge in extracting variant–function–pathogenicity associations directly from biomedical literature by leveraging the power of large language models (LLMs). Through systematic benchmarking of models, prompting strategies, and normalization pipelines, we demonstrated that a 70B instruction-tuned model, used without finetuning, can achieve high performance in extracting genetic variants and their functional annotations. With few-shot prompting, PubMind achieves >90% accuracy in variant extraction with minimal hallucination, while capturing disease terms with 99.9% precision. Importantly, the resulting annotations show strong concordance with curated resources such as ClinVar ^1^ with 100% agreement on ClinVar’s expert-reviewed variants (4 stars), yet exhibit substantial novelty, with only ∼10% overlap with ClinVar entries. By processing 41 million PubMed abstracts and 5.4 million PMC full-text articles, PubMind generated one of the most comprehensive literature-derived variant knowledgebases to date, spanning SNVs, gene fusions, SVs, and CNVs, each annotated with disease associations, pathogenicity, reasoning for pathogenicity, and source references.

Developing PubMind required addressing several key challenges: (1) accuracy of extraction, (2) efficiency of large-scale inference, (3) normalization of unstructured outputs, and (4) comparison to find undocumented variants relative to existing databases. Below we discuss each of these challenges.

### Accuracy

Benchmarking across multiple LLaMA ^15^ and DeepSeek-R1 models ^29^ confirmed that few-shot prompting and CoT reasoning provides high extraction accuracy. Cases deemed “errors” in PubMind were often formatting differences rather than true mistakes (e.g., “ΔG91” vs. “G91del”). Unlike conventional NLP approaches such as tmVar3 ^5^ or PubTator3 ^6^, which classify tokens, LLMs generate semantically consistent reformulations—often yielding clearer or standardized representations. Notably, even the base instruct model (LLaMA3.3-70B ^15^) performed well without finetuning, highlighting the potential for transferability to other biomedical information extraction tasks (e.g., drug–drug interactions, protein–protein interactions).

### Efficiency

Running LLM inference on millions of full texts is computationally infeasible. For example, processing just 500 PMC articles required ∼3.5 hours on 4 Nvidia A100 GPUs. To overcome this, we developed and evaluated two different filtering strategies: (i) regex-based filtering to reduce input volume based on key words and predefined rules and (ii) fine-tuned BERT models to retain semantically relevant paragraphs (**Supplementary Table 3**). To generate the finetuning dataset for BERT-model, we used a hybrid approach, where LLMs generate training labels for a smaller BERT classifier, enabled efficient filtering while preserving high-value content. Compared with rule-based regex filtering, fine-tuned BERT retained fewer paragraphs yet yielded more extracted variants, underscoring its semantic precision.

### Normalization

Because LLM outputs are inherently unstructured, we developed a post-processing pipeline to standardize variant, disease, phenotype, and pathogenicity annotations. Variants were parsed using regex-based normalization; diseases and phenotypes were mapped to MONDO ^21^ and HPO terms ^22^ via PubMedBERT embeddings; and pathogenicity assignments were consolidated across multiple records. Inspired by ClinVar ^1^, we further implemented a tiered confidence system to score annotation reliability, enabling transparent downstream use.

### Complementary role to existing databases

As a primary application of PubMind, we processed over 41 million PubMed abstracts and over 5.6 million full texts, to build a variant-disease-pathogenicity knowledgebase enriched with contextual information, PubMind-DB. Comparison with ClinVar ^1^ confirmed that PubMind-DB captures both reliable concordant variants and a large body of previously un-curated knowledge, with ∼90% of variants absent from ClinVar. Agreement with ClinVar improved with review depth (reaching 100% for four-star variants), supporting the validity of PubMind’s assignments. Cross comparisons with 51 computational predictors revealed strong correlation in highly pathogenic regions, while PubMind uniquely identified benign calls aligned with functional evidence but overlooked by tools. A case study on IRF6 VUS ^16^ highlighted PubMind’s ability to aggregate dispersed evidence across publications, achieving complete concordance with experimental zebrafish rescue assays—whereas AlphaMissense ^25^ and ensemble predictors ^26^ failed to identify most benign variants. The case study also underscores PubMind’s clinical relevance and ability to reclassify VUS by integrating distributed functional evidence, compared to the HGMD ^2^ which unable to provide the pathogenicity for these variants. A second case study on PDGFRB in infantile myofibromatosis ^27, 30–32^ further illustrated how PubMind-DB not only retrieves variant–disease associations across scattered publications but also surfaces therapeutic insights, such as imatinib response, directly from the literature. Together, these case studies demonstrate PubMind’s ability to integrate dispersed evidence across the literature to accurately reclassify variants of uncertain significance and to contextualize rare disease variants with clinically actionable insights, complementing existing databases and computational predictors.

Despite these strengths, some limitations remain. First, due to computational constraints of large models (e.g., requiring multiple H100/A100 GPUs), PubMind processes abstracts and paragraphs independently rather than entire full texts. While this reduces input size and improves efficiency, it may miss cross-paragraph associations within a single study. Second, as with any generative model, outputs may deviate from the requested format; extensive normalization mitigates this but does not eliminate rare inconsistencies (e.g., cDNA vs. protein-level misassignments). Nevertheless, hallucinations were minimal, and overall extraction accuracy was consistently high.

In contrast to HGMD ^2^, which provides only variant–disease associations, and ClinVar ^1^, which depends on manual submissions and variable curation depth, PubMind offers scalable, literature-grounded extraction of explicit variant–disease–pathogenicity relationships. The pipeline can process the entire PubMed and PMC Open Access corpus in 1-2 weeks with 4 Nvidia A100 GPUs, delivering up-to-date annotations with direct links to source text. Beyond variant interpretation, the framework is generalizable to other biomedical entity–relationship extraction tasks, from pharmacogenomics to pathway annotation. Additionally, we evaluated the applicability of PubMind to in-house clinical notes for patients with known positive genetic diagnosis. In this setting, the PubMind system successfully extracted variants along with associated diseases and LLM-derived reasoning (e.g., “the gene panel confirmed the variant’s pathogenicity”). PubMind was able to efficiently filter through dozens of clinical notes for a single patient, retaining only those relevant for variant extraction. The resulting structured outputs provide a rapid, high-level index of patient data, offering a valuable foundation for clinical data management and downstream analysis.

In summary, PubMind represents a powerful and novel bridge between unstructured biomedical text and structured genomic resources. By integrating advanced LLM reasoning with efficient filtering and rigorous normalization, PubMind delivers both scale and interpretability, complementing curated databases and computational predictors. We anticipate that PubMind and PubMind-DB will serve as valuable pipeline and resource for human variant interpretation, evidence-based variant reclassification, rare disease gene discovery, and precision medicine applications.

## Materials and Methods

### Data sources and corpus

For the establishment of PubMind-DB, 41,682,357 PubMed Abstracts and 5,425,084 PubMed Central (PMC) full texts from OA subset have been downloaded (up to 2025.02.07) through the NCBI’s ftp depository (https://pubmed.ncbi.nlm.nih.gov/download/, https://pmc.ncbi.nlm.nih.gov/tools/ftp/). XML files were parsed using the Python package BeautifulSoup4 to extract PubMed abstracts or PMC full texts.

### Input filtering for PubMed abstracts and PMC full texts

We prepared different databases for finetuning and benchmarked different BERT models for the task of input filtering (**Supplementary Table 3**). To filter the enormous literature input, DistilBERT model (https://huggingface.co/distilbert/distilbert-base-uncased) was finetuned using 1.5K labelled abstracts. The finetuned DistilBERT classified each paragraph according to whether it contained variant information. The 1.5K labeled dataset consisted of ∼1000 negative examples generated using regex rules and 500 positive examples from the PubTator3.0 training corpus ^6^. Additional BERT models have been used and tested, including GoogleBERT ^13^, BioMedBERT ^33^. Larger finetuning datasets of 15K abstracts were generated and evaluated using two labeling strategies: regex-based labels (regex-label) and LLM-derived labels (LLM-label). For LLM-labels, we first ran 100,000 PubMed abstracts through the LLM; abstracts producing useful variant outputs were labeled positive, while those without were labeled negative. See **Supplementary Methods** for details of finetuning dataset construction.

### Model selection and prompt engineering for LLM inference

To evaluate whether prompt engineering with base LLM models (without finetuning) could enable genetic variant extraction, we experimented different prompts and compared the results with benchmark dataset based on 500 PubMed abstracts with labelled variant information ^6^. Different versions of LLaMA family ^15^ and DeepSeek-R1 models ^29^ were evaluated (**Supplementary Figure 1**). After choosing LLaMA3.3-70B instruct model as our main model, different prompts were evaluated based on the accuracy of variant extraction and the GPU runtime (**Supplementary Figure 2**). For details about all the prompts we used, please refer to **Supplementary Table 1**.

### Variant and disease extraction benchmark

To assess reliability and accuracy, we benchmarked the LLM’s ability to extract both variants and diseases. The benchmark dataset for variant NER is the 500 PubMed abstracts with labelled variant from PubTator3 ^6^. The benchmark dataset for disease NER is from NCBI disease corpus ^34^ used in PhenoTagger ^35^, which contains the PubMed abstract as well as the disease name found in the text. Hallucination rates were quantified by comparing all LLM outputs against these benchmark datasets (**Supplementary Table 4&5**).

### Normalization and Database Generation

The variants (RSID, cDNA change, amino acid change) were normalized using regular expression (regex), and the transcript information and corresponding genomic coordinates for variants were extracted using PyEnsembl python package. The LLM output disease was normalized using MONDO human disease name ^21^, while the LLM output phenotype was normalized using HPO term ^22^. The normalization is based on the cosine similarity >0.9 using PubMedBERT embedding (NeuML/pubmedbert-base-embeddings from hugging face) for MONDO disease name and HPO term compared with LLM output disease and phenotype. The gene names were filtered based on Ensembl v111 ^36^ to remove non-human genes from literatures. The final PubMind pathogenicity has been consolidated using all records from different sources of publication. The PubMind pathogenicity score is calculated based on the equation below:

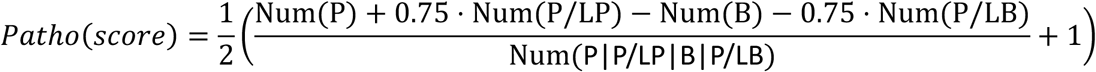

where P=pathogenic, LP= Likely Pathogenic, B=Benign, LB=Likely Benign.

### Pathogenicity annotation and assessment

To annotate the variants with genome coordinates, ANNOVAR ^23^ was used to perform variant annotation, for both gene and functional based, population-based, and bioinformatics tools-based analyses. For gene and functional annotation, hg38 human genome from Ensembl ^36^ (2024-05-13) was used. For population-based annotation, gnomAD ^37^ (version 4.1 whole-genome data) and ClinVar ^1^ (date: 20240917) database was used to cross check the population allele frequency and pathogenicity summarization. For bioinformatics pathogenicity predictions, dbNSFP version 4.7a ^24^ was used to get pathogenicity predictions from 23 bioinformatics software, including AlphaMissense ^25^ and MetaRNN ^38^.

### Data and website accessibility

The PubMind source code is available at GitHub (https://github.com/WGLab/PubMind). The PubMind knowledgebase is accessible via a web interface (http://PubMind.wglab.org/) and API. The web application was built with Flask (Python) and supports data queries using SQLite3.

## Data Availability

PubMed Abstracts and PubMed Central (PMC) full texts can be accessed through the open-access ftp depository (https://pubmed.ncbi.nlm.nih.gov/download/, https://pmc.ncbi.nlm.nih.gov/tools/ftp/).

## Code Availability

PubMind is released under open-source license and can be accessed here: https://github.com/WGLab/PubMind. The PubMind-DB can be accessed here: http://pubmind.wglab.org/.

## Supplementary Information

Supplementary Materials. A Word document contains supplementary methods, tables and figures.

