## Supplementary Materials for "PubMind: Literature-Based Genetic Variant Extraction and Functional Annotation Using Large Language Models"

### Supplementary Methods

#### Prompt engineering

We initial tested different version and size of Llama models, and Llama3.3-70b had the best overall performance (**Supplementary Figure 1**). Moving further with Llama3.3-70b, we found that the biggest influence of the performance of genetic variant NER is the user prompt (**Supplementary Figure 2**). Few shots prompt (few-shot) and Chain of Thought (CoT) prompt showed higher accuracy compared to zero-shot (0-shot). The order of the input paragraph of literature also influenced the results in CoT prompt, which showed that putting paragraph after CoT prompt had higher accuracy in cDNA variant extraction, which might be caused by the forgetting of user instruction after a long input paragraph. The system prompt did not play much effect here in genetic variant extraction (see **Supplementary Table 1** for all the prompts). Overall, a short system prompt with few-shot example showed the most consistent and reliable accuracy (near 90%), as well as efficient LLM inference time, for both cDNA mutation and protein mutation. We also checked the 10% of the LLM-extracted variant for why they are not the same as the benchmarked dataset, many of them are actually correct variants but they have different expression (i.e., ‘A>G at nucleotide 313’ in label and ‘A313G’ in LLM output). Most of the difference is from the fundamental difference of a BERT-based output and GPT-based approach, where the label for a BERT model is classification of the original tokens, but GPT is a new generated token based on the summarization of the GPT model (check **Supplementary Table 4** for detail). Therefore, the based LLM model (Llama3.3 70B) showed considerable few-shot ability in terms of genetic variant NER without any finetuning.

#### Input filtering techniques and training dataset construction

We implemented two filtering strategies: rule-based regex filtering and BERT-based filtering. For PubMed abstracts, filtering was applied at the abstract level; for PMC full texts, it was performed paragraph by paragraph. To generate training data for BERT finetuning, we constructed three datasets: (1) 1.5k dataset (regex-label): 500 positive abstracts from the PubTator3.0 corpus ^1^ and 1,000 regex-labeled negatives. (2) 15k dataset (LLM-label): 10,000 positives and 5,000 negatives labeled based on LLM output (abstracts producing useful variant outputs vs. not). (3) 15k dataset (hybrid): labels consolidated from regex-label and LLM-label.

As shown in **Supplementary Table 3**, the fine-tuned BERT models produced more enriched variant outputs while filtering out a larger fraction of irrelevant text, demonstrating the advantage of BERT-based input filtering. Among training datasets, the simplest regex-based 1.5k dataset achieved the best efficiency: it retained fewer paragraphs overall but yielded the most useful LLM outputs. While BioMedBERT achieved the highest filtering efficiency, the final quality of LLM outputs was lower than that obtained with DistilBERT. Considering both efficiency and output quality, DistilBERT was selected as the primary fine-tuned model for raw input filtering.

#### Normalization of variant, disease, phenotype, and pathogenicity

After the benchmarking of variant extraction, disease recognition, and LLM hallucination, the instruct-tuned Llama3.3-70B base model demonstrated a reasonable and confident NER suitable for our variant-disease association extraction task. Our next step is to make sure the variant, disease and phenotype extracted by LLM is standardized and easily transferable to other widely-used formats. To complete the tasks, both regex-based and cosine similarity-based approaches were used. For variant extraction, specifically for single nucleotide variants (SNVs), we used a complex regex to confirm the cDNA change and amino acid change based on LLM extracted DNA mutation and protein mutation. The SNVs that passed our regex rules will be considered a reasonable SNV and will be further normalized using Ensembl v111 ^2^ to extract all possible transcripts for a cDNA/protein change and provide all possible genome coordinates for the mutation.

We calculated the cosine similarity of LLM-output disease and phenotype with MONDO human disease name and HPO term correspondingly, based on both PubMedBERT embedding and string similarity (**Supplementary Fig 3a-b)**. Based on the distribution, we can see a relative normal distribution for the similarity of LLM-output phenotype with HPO term, but a very distinct profile with a peak at 1 for disease similarity. The different distributions of phenotype and disease showed that the disease name LLM extracted from the literature is more standardized compared to phenotype, hence a distinct distribution is observed.

For disease outputted by LLM, we used MONDO disease name for human (n=22,758) and a 0.9 cosine similarity based on PubMedBERT embedding as a cutoff. For phenotype outputted by LLM, HPO term (n=19,533) and 0.9 cosine similarity was used to keep high confident phenotype. After the normalization, we will have a high confident genetic variant knowledgebase which contains high confident SNV with VCF-format genome coordinates, as well as corresponding MONDO disease and HPO phenotype extracted from the original literature text.

#### Benchmark and hallucination evaluation for disease extraction

To further evaluate the ability of disease recognition using instruction-tuned LLaMA3.3-70B model, we performed a benchmark using 593 PubMed abstracts with labelled diseases. To compare the accuracy of LLM extracted disease with ground truth disease in NCBI corpus, we calculated the cosine similarly for the PubMedBERT embedding (see **Method**) of LLM-output disease and ground truth disease. Based on the **Supplementary Figure 3c**, there are 647 LLM extracted diseases with > 0.8 cosine similarity with the ground truth, and 82 disease with < 0.8 cosine similarity are further investigated. Among the 82 LLM inferred disease name, 65 of them found exact string matched in the original abstracts, and 17 of them were not exactly the same as original input but they are similar and valid alternative expressions (see **Supplementary Table 5** for details). There is only 1 LLM output disease “breast neoplasms” corresponding to “breast and squamous cell neoplasms” in the original abstract, which might be considered somewhat inaccurate. Therefore, the based LLaMA3.3-70B instruct model with appropriate few-shot prompts demonstrated the >0.999 accuracy (728/729) and 0 hallucination in terms of disease name recognition.

#### Variant Consolidation

In PubMind, each variant is uniquely identified based on the PubMind Variant ID (PVID). Each PVID is a unique variant and is an aggregation of multiple records from separated LLM output. The SNVs will be the most standardized variant type and it will be identified by gene name together with either RSID, or cDNA change or protein change. For the SNVs that match the Ensembl transcript, there will be genomic coordinates available in VCF-format. A SNV will have disease and phenotype information associated with the record, as well as transcript and genome coordinates information if available.

To consolidate the pathogenicity assignment for each variant that sometimes contain multiple pathogenicity from different paragraph or literatures. Similar approach like ClinVar was used, which summarize the final pathogenicity (pathogenicity_sum) into these 6 categories: Benign, Benign/Likely Benign (B/LB), Pathogenic, Pathogenic/Likely Pathogenic (P/LP), Conflicting, and Uncertain significance. The conflicting of pathogenicity means that there are contradict pathogenicity from different records of the same variant, which could come from different paragraph of the same paper or from different papers. The unknown pathogenicity means there is neither benign/likely benign assignment nor pathogenic/likely pathogenic assignment for the variant. The pathogenicity score is calculated based on number of pathogenic, P/LP, benign, B/LB over the total number of the total records with pathogenicity assignment (see **Method** for detail), with 1 represents the most pathogenic and 0 represents the most benign.

### Supplementary Tables and Figures

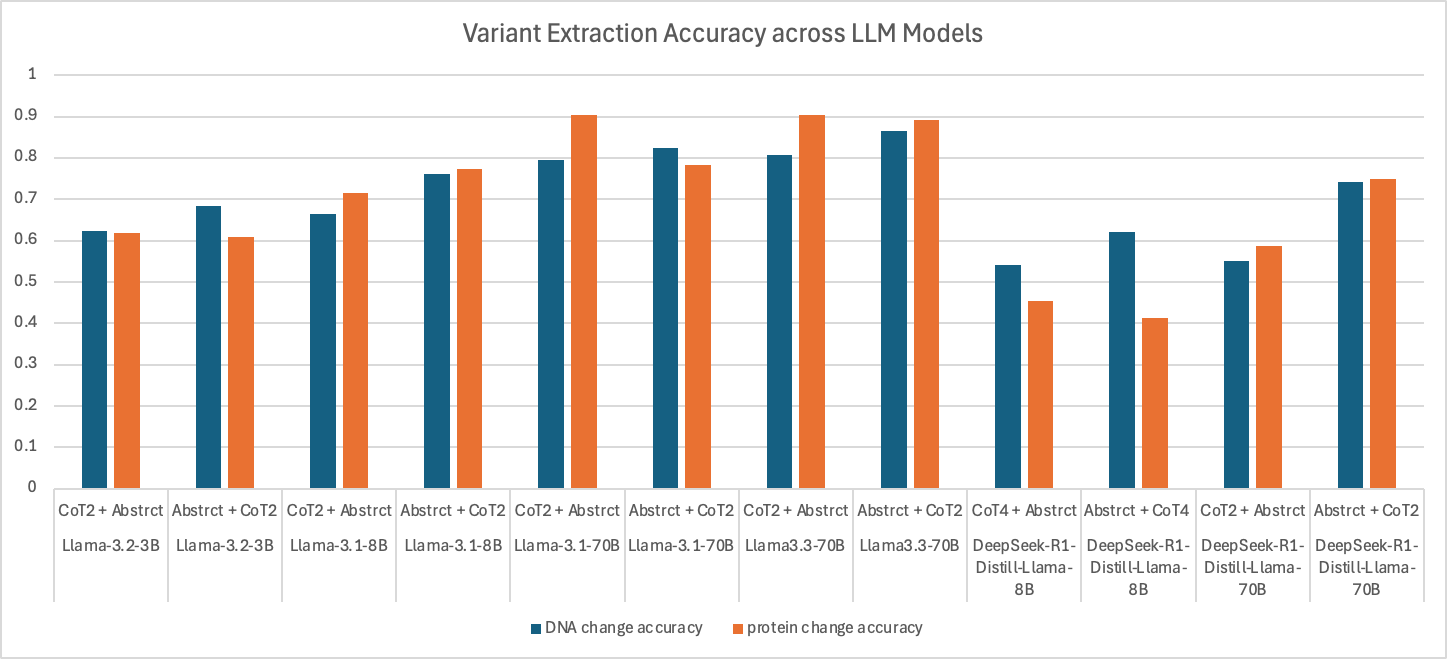

**Supplementary Figure 1**. The variant extraction benchmark using different LLaMa3 family and DeepSeek-R1 models, with different order of prompt (CoT2) and abstract. Overall, The LlaMA3.3-70b model showed the best performance for both DNA change extraction and protein change extraction. Please refer to **Supplementary Table 1** for details about the CoT2.

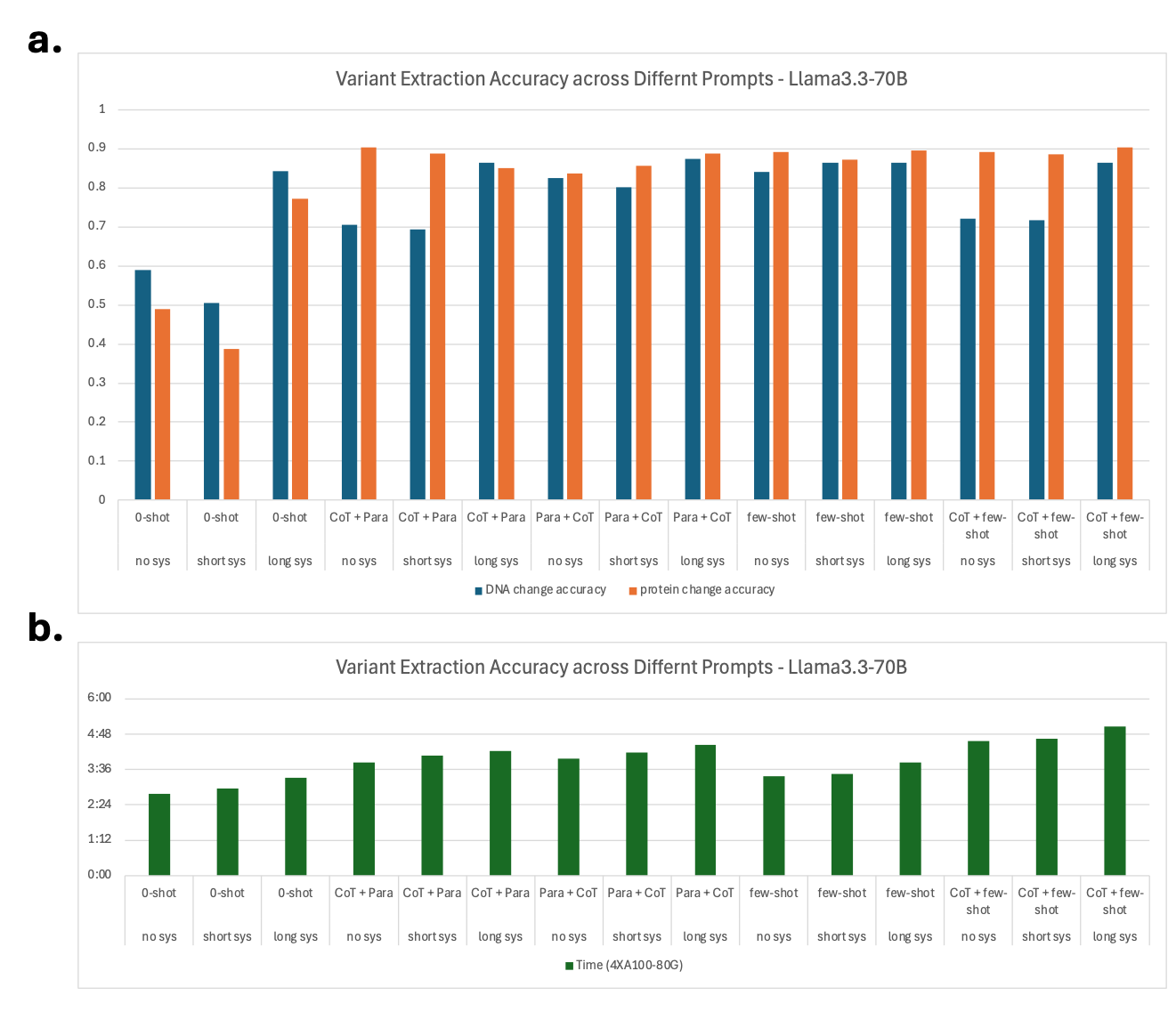

**Supplementary Figure 2**. Impact of system and user prompts on variant extraction accuracy using LLaMA3.3-70B. (a) Accuracy of variant extraction under different prompting strategies. Bars show DNA change accuracy (blue) and protein change accuracy (orange). Prompting conditions tested included zero-shot (0-shot), Chain-of-Thought (CoT), paragraph-ordering (Para), few-shot examples, and combinations thereof. The order of prompts and paragraph placement was also evaluated. (b) GPU runtime (mm:ss) for each prompting strategy, measured on 4× NVIDIA A100-80G GPUs. While few-shot and CoT prompts yielded the highest extraction accuracy (~90%), they required longer inference times compared to zero-shot prompting.

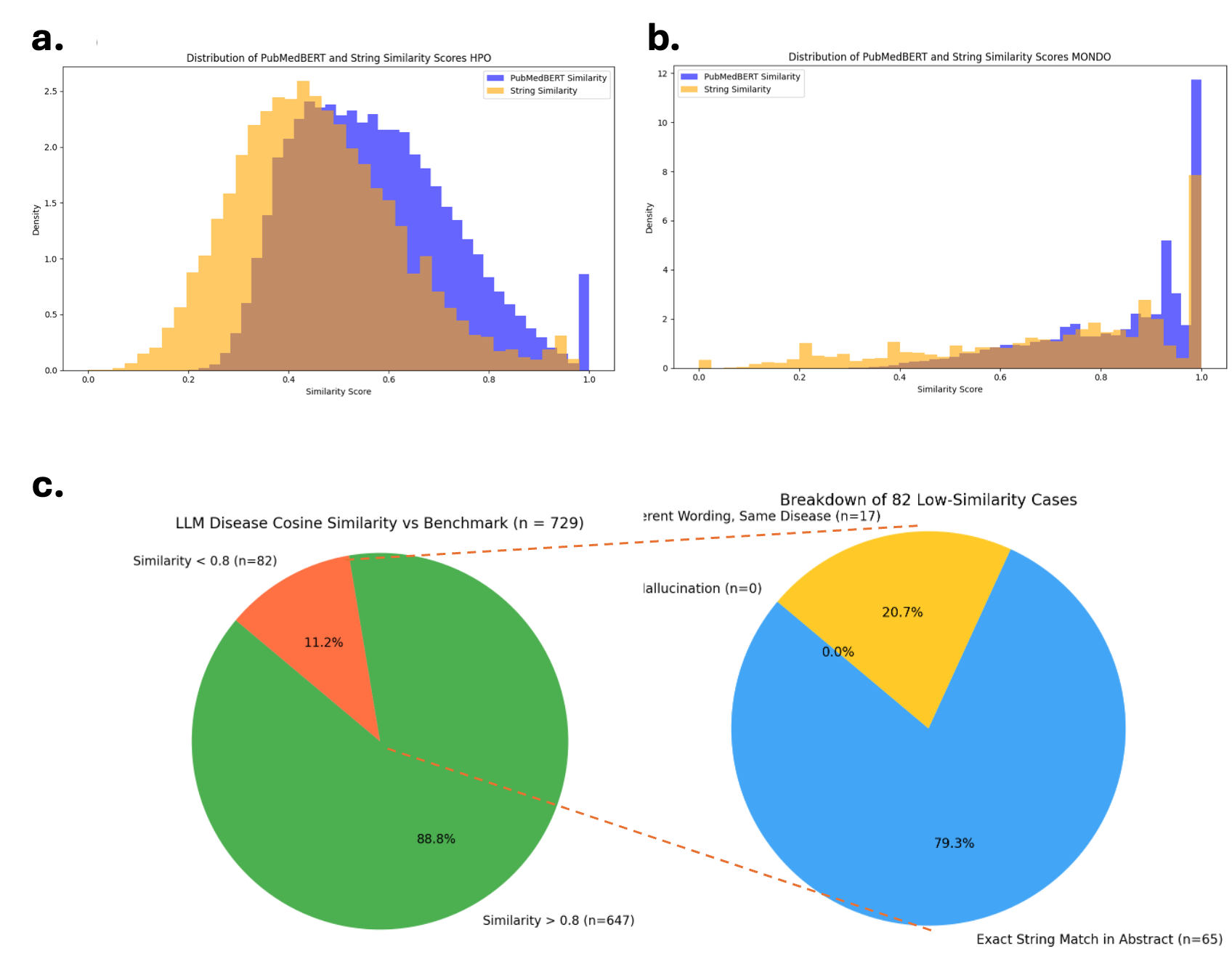

**Supplementary Figure 3**. Evaluation of normalization and disease extraction accuracy. (a–b) Distribution of similarity scores for LLM-extracted terms compared against standard ontologies. PubMedBERT cosine similarity (blue) and string similarity (orange) are shown for (a) HPO and (b) MONDO. Disease terms display a sharp peak at 1.0, reflecting standardized wording, while phenotypes show a broader distribution due to more variable expression. (c) Disease NER benchmarking. Comparison of LLM-extracted diseases (n=729) against the NCBI disease corpus. Left pie chart: 88.8% of predictions achieved cosine similarity >0.8, while 11.2% fell below. Right pie chart: of the 82 low-similarity cases, 65 (79.3%) were exact string matches in the abstracts, 17 (20.7%) were valid alternative expressions, and none represented hallucinations.

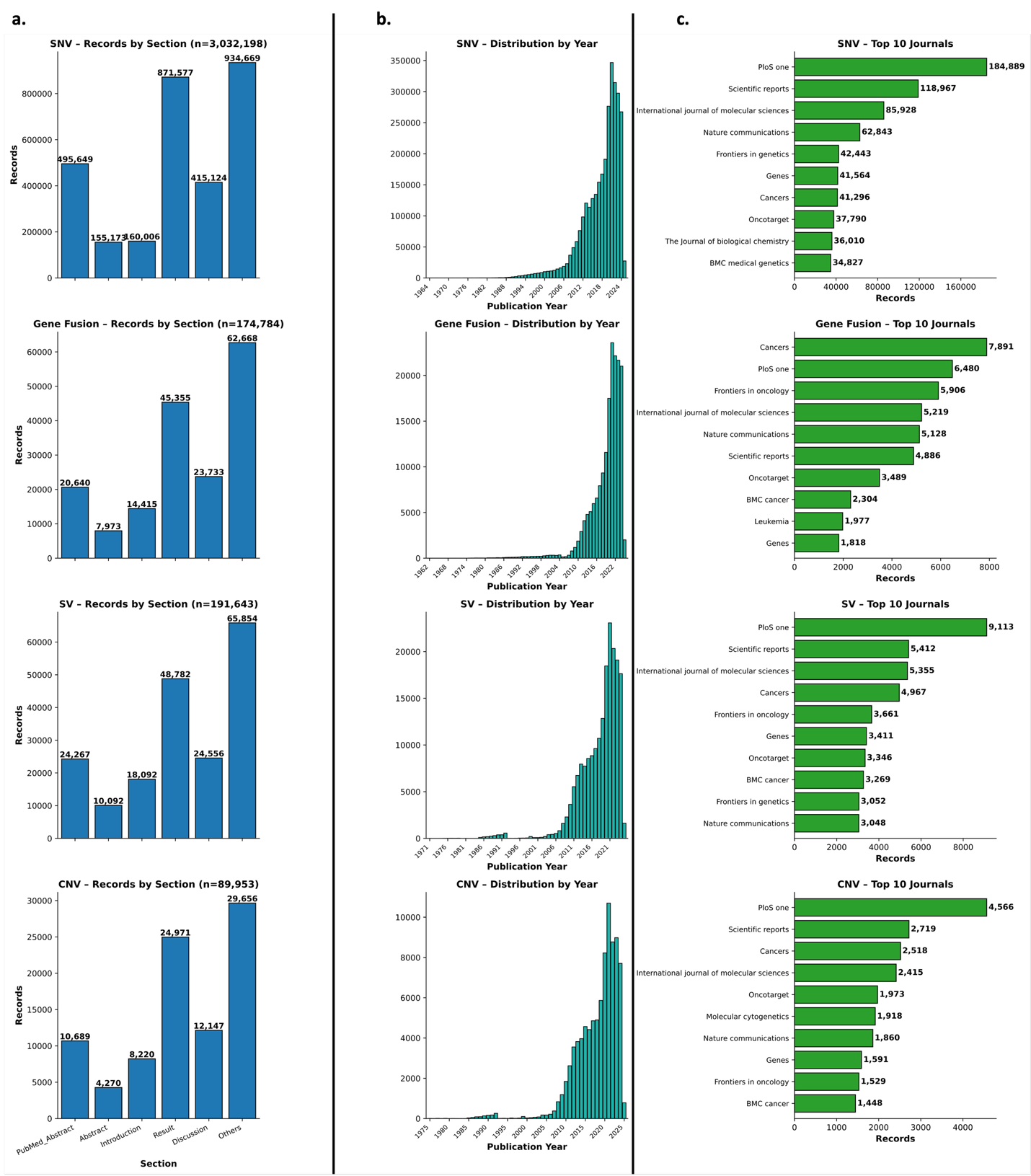

**Supplementary Figure 4**. Distribution of PubMind variant records by literature section, publication year, and journal. (a) Section-wise distribution. Variant records extracted from PubMed abstracts and PMC full texts, stratified by document section, for SNVs (n=3,032,198), gene fusions (n=174,784), SVs (n=191,643), and CNVs (n=89,953). Most variants were identified in sections labeled “Other,” which include XML fields not mapped to standard categories. (b) Temporal distribution. Publication year of extracted variants across all four variant types, showing steady growth with a sharp peak in 2021. (c) Journal distribution. Top ten journals contributing variant records for each variant type. PLOS One and Scientific Reports consistently rank among the highest, reflecting their large volume of genetics publications.

**Supplementary Table 1**. All user prompts and system prompts used for the variant extraction benchmark.

| **User Prompt** | |
| --- | --- |
| 0-shot | """  Extract the information of gene, DNA mutation, protein mutation, related disease, experiment result, and pathogenicity from this research paper paragraph in the format of ##gene, DNA mutation, protein mutation, disease, experimental result, pathogenicity.  """ |
| Few-shot | """  Extract the information of gene, DNA mutation, protein mutation, related disease, experiment result, and pathogenicity from this research paper paragraph. Answer exactly in the format of ##gene, DNA mutation, protein mutation, disease, experimental result, pathogenicity.  Example 1:  Paragraph: "Paragraph: For ESRP2, variants R250Q, R353Q and R667C rescued the molecular splicing of Arhgef11 in the Py2T assay, (Fig. 3E, Table 1)."  Output:  ##ESRP2, -, R250Q, orofacial cleft (OFC), rescued the molecular splicing of Arhgef11 in the Py2T assay, benign  ##ESRP2, -, R353Q, orofacial cleft (OFC), rescued the molecular splicing of Arhgef11 in the Py2T assay, benign  ##ESRP2, -, R667C, orofacial cleft (OFC), rescued the molecular splicing of Arhgef11 in the Py2T assay, benign  Example 2:  Paragraph: "The mutation c.123A>T in BRCA1 leads to a truncated protein p.Lys41*, associated with breast cancer. Functional assays showed loss of DNA repair function, indicating pathogenicity."  Output:  ##BRCA1, c.123A>T, p.Lys41*, breast cancer, loss of DNA repair function, pathogenic  Example 3:  Paragraph: "A variant in CFTR, p.Phe508del, causes cystic fibrosis. Studies confirmed defective chloride channels, confirming pathogenicity."  Output:  ##CFTR, -, p.Phe508del, cystic fibrosis, defective chloride channels, pathogenic  """ |
| CoT (for Sup Fig 2) | """  Extract the information of gene, DNA mutation, protein mutation, related disease, experiment result, and pathogenicity from this research paper paragraph in the format of ##gene, DNA mutation, protein mutation, disease, experimental result, pathogenicity.  Follow these steps to ensure accuracy:  1. Identify Relevant Genes and Mutations, locate any gene names or symbols mentioned in the paper.  2. Identify if there is any DNA-level mutation. Make sure the DNA mutation only contains letter A,T,C,G. Note any variant start with 'c.'.  3. Identify if there is any protein-level mutation. Note different expression of a protein or amino acid, no matter in three letter or one letter. If only an RSID is found, fill it in as the DNA mutation. Note any variant start with 'p.'.  4. Determine Disease Association: identify any diseases or conditions related to each gene and mutation. Look for terms like "disease," "syndrome," "condition," or names of specific conditions.  5. Extract Functional or Experimental Result: find statements describing the effect of each mutation, especially in terms of experimental assays or functional impacts. Note results like "rescued" or "failed to rescue" a function.  6. Assign Pathogenicity Level: determine if the mutation is "pathogenic," "benign," or similar terms indicating pathogenicity or clinical relevance.  7. Format Output Concisely: present the result in this format: ##gene, DNA mutation, protein mutation, disease, experimental result, pathogenicity.  Using these steps, extract and present the unique ##gene, DNA mutation, protein mutation, disease, experiment result, pathogenicity for each mutation-disease relationship in this research paper paragraph/abstract:  """ |
| CoT2 (for Sup Fig 1) | """  Extract the information of gene, DNA mutation, protein mutation, related disease, experiment result, and pathogenicity from this research paper paragraph in the format of ##gene, DNA mutation, protein mutation, disease, experimental result, pathogenicity. Follow these steps to ensure accuracy:  1. Identify Relevant Genes and Mutations, locate any gene names or symbols mentioned in the paper.  2. Identify if there is any DNA-level mutation. Make sure the DNA mutation only contains letter A,T,C,G. Note any variant start with 'c.'.  3. Identify if there is any protein-level mutation. Note different expression of a protein or amino acid, no matter in three letter or one letter. If only a RSID is found, fill it in as the DNA mutation. Note any variant start with 'p.'.  4. Determine Disease Association: identify any diseases or conditions related to each gene and mutation. Look for terms like "disease," "syndrome," "condition," or names of specific conditions.  5. Extract Functional or Experimental Result: find statements describing the effect of each mutation, especially in terms of experimental assays or functional impacts. Note results like "rescued" or "failed to rescue" a function.  6. Assign Pathogenicity Level: determine if the mutation is "pathogenic," "benign," or similar terms indicating pathogenicity or clinical relevance.  7. Format Output Concisely: present the result in this format: ##gene, DNA mutation, protein mutation, disease, experimental result, pathogenicity  Using these steps, extract and present the unique ##gene, DNA mutation, protein mutation, disease, experiment result, pathogenicity for each mutation-disease relationship in this research paper only once. Stop respond when you find all mutations.  """ |
| Gene Fusion few-shot prompt | """Extract gene fusion information from the paragraph in the format:  ##gene fusion\|\|driver gene\|\|partner gene\|\|domain affected\|\|related disease\|\|experiment or functional result\|\|pathogenicity  - The **gene fusion** should be in the format **GENE1::GENE2** as mentioned in the text.  - Identify the **driver gene** — this is typically the gene contributing the active domain (e.g., kinase, transcription factor) or the 3' gene in oncogenic fusions. Use context clues (e.g., "drives", "activates", "encodes") to determine.  - The **partner gene** is the fusion partner that contributes less functional consequence or is upstream/regulatory.  Example 1:  "The EML4-ALK fusion occurs in lung cancer and activates the ALK kinase domain, leading to cell proliferation in vitro."  ##EML4::ALK\|\|EML4\|\|ALK\|\|kinase domain\|\|lung cancer\|\|cell proliferation in vitro\|\|pathogenic  Example 2:  "A rare fusion between TMPRSS2 and ERG was found in prostate cancer, with no functional assay performed."  ##TMPRSS2::ERG\|\|TMPRSS2\|\|ERG\|\|-\|\|prostate cancer\|\|not tested\|\|unknown  Example 3:  "We identified a novel RUNX1-RUNX1T1 fusion in acute myeloid leukemia, disrupting the DNA-binding domain of RUNX1."  ##RUNX1::RUNX1T1\|\|RUNX1\|\|RUNX1T1\|\|DNA-binding domain\|\|acute myeloid leukemia\|\|disrupts DNA binding\|\|pathogenic  Now extract from the paragraph below:  """ |
| CNV few-shot prompt | """  You are given a biomedical paragraph. Extract any copy number variants (CNVs) mentioned, including deletions or duplications. For each CNV, extract the following structured fields:  ##CNV_type\|\|chromosome_region\|\|gene(s)\|\|genomic_coordinates\|\|disease\|\|experiment or functional result\|\|pathogenicity  - CNV_type: "deletion" or "duplication"  - chromosome_region: cytogenetic band (e.g., 22q11.2) if available  - gene(s): affected gene(s), separated by commas if multiple  - genomic_coordinates: if available (e.g., chr22:18000000-21000000)  - disease: associated disease name, or "unknown"  - experiment or functional result: brief summary of experimental or clinical findings  - pathogenicity: must be one of ["pathogenic", "likely pathogenic", "benign", "likely benign", "unknown", "conflicting"]  Example 1:  "The 22q11.2 deletion syndrome is caused by a ~3Mb deletion on chromosome 22, encompassing the TBX1 gene, and is associated with DiGeorge syndrome."  Answer:  ##deletion\|\|22q11.2\|\|TBX1\|\|chr22:18000000-21000000\|\|DiGeorge syndrome\|\|3Mb deletion spanning TBX1\|\|pathogenic  Example 2:  "A microduplication of 16p11.2 involving the SH2B1 gene was observed in individuals with early-onset obesity."  Answer:  ##duplication\|\|16p11.2\|\|SH2B1\|\|unknown\|\|early-onset obesity\|\|Reported in multiple obesity patients\|\|likely pathogenic  Example 3:  "An inherited 1q21.1 deletion encompassing the GJA5 gene was identified in a family with variable phenotypes, but with no consistent clinical presentation."  Answer:  ##deletion\|\|1q21.1\|\|GJA5\|\|unknown\|\|unknown\|\|Detected in unaffected individuals and patients\|\|conflicting  Example 4:  "A 2q13 deletion involving NPHP1 was found in both patients with kidney disease and unaffected family members."  Answer:  ##deletion\|\|2q13\|\|NPHP1\|\|unknown\|\|kidney disease\|\|Lack of consistent phenotype among carriers\|\|likely benign  Now extract the CNV(s) from the following paragraph:  """ |
| SV few-shot prompt | """  You are given a biomedical paragraph. Extract any **structural variants (SVs)** mentioned, including deletions, duplications, inversions, insertions, and translocations. For each SV, extract the following structured fields:  ##SV_type\|\|gene(s)\|\|chromosome_region\|\|genomic_coordinates\|\|related disease\|\|experimental or clinical evidence\|\|pathogenicity  - SV_type: one of ["deletion", "duplication", "inversion", "insertion", "translocation"]  - gene(s): affected gene(s), separated by commas if multiple  - chromosome_region: cytogenetic region (e.g., 22q11.2), or "unknown"  - genomic_coordinates: format "chr:start-end" if available, or "unknown"  - related disease: disease associated with the SV, or "unknown"  - experimental or clinical evidence: brief summary of supporting data  - pathogenicity: one of ["pathogenic", "likely pathogenic", "benign", "likely benign", "unknown", "conflicting"]  Example 1:  "A pathogenic inversion disrupting the F8 gene on Xq28 is a common cause of severe hemophilia A."  Answer:  ##inversion\|\|F8\|\|Xq28\|\|chrX:154180000-154300000\|\|hemophilia A\|\|Breakpoints identified in patients with severe phenotype\|\|pathogenic  Example 2:  "A translocation between chromosomes 9 and 22 creates the BCR-ABL1 fusion gene, known as the Philadelphia chromosome, found in chronic myeloid leukemia."  Answer:  ##translocation\|\|BCR,ABL1\|\|9q34,22q11\|\|chr9:133729000-chr22:23632600\|\|chronic myeloid leukemia\|\|Philadelphia chromosome detected by FISH\|\|pathogenic  Example 3:  "A 5q13.3 duplication involving the SMN1 and SMN2 genes has been reported in some individuals without symptoms."  Answer:  ##duplication\|\|SMN1,SMN2\|\|5q13.3\|\|chr5:70000000-70200000\|\|unknown\|\|Identified in asymptomatic carriers during screening\|\|benign  Example 4:  "An insertion of ~1.5 kb in intron 1 of the FGFR2 gene has been observed in individuals with Apert syndrome."  Answer:  ##insertion\|\|FGFR2\|\|10q26\|\|chr10:123800000\|\|Apert syndrome\|\|Insertion disrupts regulatory elements in FGFR2 intron 1\|\|likely pathogenic  Now extract the SV(s) from the following paragraph:  """ |
| **System Prompt** | |
| short | '''You are an expert in genomics and bioinformatics. Extract the variant-related information from scientific publication paragraphs.''' |
| long | '''  You are an expert in genomics and bioinformatics. Extract the following from scientific publication paragraphs:  - **Gene names** (official symbols).  - **DNA mutations** (A, T, C, G, or starting with 'c.').  - **Protein mutations** (amino acid changes, e.g., 'p.' notation).  - **Associated diseases**.  - **Experimental results** (e.g., "loss of function").  - **Pathogenicity** (e.g., "pathogenic," "benign").  Output in this format:  `##gene, DNA mutation, protein mutation, disease, experimental result, pathogenicity`  Use a dash (`-`) if data is missing. Only extract what is explicitly mentioned—no assumptions.  ''' |

**Supplementary Table 2**. Criteria for assigning PubMind confidence tiers. The PubMind confidence score reflects the reliability of each variant annotation, based on literature support, consistency of pathogenicity, and availability of genomic mapping. Each satisfied rule contributes +1 to the overall confidence score.

| **Rule** | **Confidence criteria** | **Abbreviation in PubMind** |
| --- | --- | --- |
| 1 | Supported by multiple publications (Num_of_paper_used > 1) | multiple papers |
| 2 | Pathogenicity assignments are consistent across records (not “Conflicting” or “Uncertain significance”) | consistent pathogenicity |
| 3 | Variant is mapped to genomic coordinates (transcript identified) | transcript found |

**Supplementary Table 3**. The comparison of raw literature filtering using different method (regex vs. BERT model) as well as training datasets. The efficiency is calculated based on the total number of high quality LLM output of genetic variants divided by the total number of filtered paragraph as input.

| **Model & finetune dataset** | **Model Size (Num of parameters)** | **Raw Paragraph** | **Final filtered input** | **LLM raw output** | **LLM output** | | **LLM quality output (pass quality control)** | **raw output / input** | **quality output / raw output** | **Overall Efficiency (quality output / input)** |
| --- | --- | --- | --- | --- | --- | --- | --- | --- | --- | --- |
| regex filter | N/A | 826393 | 14804 | 3745 | 2743 | 2203 | | 0.25297217 | 0.73244326 | 0.18528776 |
| **BioMedBERT + 1.5k_dataset** | **109483778** | **826393** | **6307** | **14735** | **3987** | **3005** | | 2.33629301 | 0.27058025 | 0.63215475 |
| **distillBERT + 1.5k_dataset** | **66955010** | **826393** | **10326** | **23559** | **4565** | **3260** | | 2.28152237 | 0.19376884 | 0.44208793 |
| **googleBERT_cased** ^3^ **+ 1.5k dataset** | **108311810** | **826393** | **12080** | **22464** | **4427** | **3177** | | 1.85960265 | 0.19707087 | 0.36647351 |
| distillBERT + 15k_LLM_regex_label | 66955010 | 826393 | 12107 | 22454 | 4418 | 3167 | | 1.85462955 | 0.19675782 | 0.36491286 |
| googleBERT_cased + 15k_LLM_label_only | 108311810 | 826393 | 23608 | 46892 | 5626 | 3623 | | 1.98627584 | 0.11997782 | 0.23830905 |
| BioMedBERT ^4^+ 15k_LLM_label_only | 109483778 | 826393 | 38396 | 61854 | 6414 | 3887 | | 1.61094906 | 0.1036958 | 0.16704865 |

**Supplementary Table 4**. Some examples of the DNA change and protein change difference of the PubTator3 training label with LLM output. The label is based on the PubTator3 train and test dataset.

| **PMID** | **Type of mutation** | **Label** | **LLM output** | **Num of record in label** | **Num if record in LLM output** | **Num of match record** | **Difference (match - label)** | **Comment** |
| --- | --- | --- | --- | --- | --- | --- | --- | --- |
| 19880293 | DNA | {'6054,deletion/ATT', '3279,A/C', '6054 deletion/ATT', '924,A/G', 'IVS9+459,A/G', '924A/G'} | {'3279A/C', '6054deletion/ATT', 'IVS9+459A/G', '924A/G'} | 6 | 4 | 2 | 4 | 6054,deletion/ATT' and '6054deletion/ATT' are the same mutation, LLM just output one of them; same for '924,A/G' and '924A/G' |
| 15316799 | DNA | {'intron2,352G>A', 'intron4,+718C>T', '15T>C', 'intron3,+45C>T'} | {'352G>A', '+718C>T', '+45C>T', '15T>C'} | 4 | 4 | 1 | 3 | LLM did not include intron number, which is not necessary if position is provided |
| 16252083 | DNA | {'A>G transition at 127 position', 'G>A transition at 1032 position'} | {'G1032A', 'A127G'} | 2 | 2 | 0 | 2 | LLM summarized the unstructed text into well-formmated variant |
| 20887110 | DNA | {'C>T transition at nucleotide 677', '677C>T', 'C677T'} | {'C677T'} | 3 | 1 | 1 | 2 | All three expressions in the label are the same, LLM just output one |
| 19429592 | protein | {'E299sp', 'G289sp', 'S522fs525sp', 'E169sp'} | {'G289*', 'S522fs*25', 'E299*', 'E169*'} | 4 | 4 | 0 | 4 | LLM use * instead of "sp" to represent a stop, which is more accurate |
| 16252083 | protein | {'N>S substitution at codon 127', 'G322D', 'G>D change at codon 322', 'N127S'} | {'N127S', 'G322D'} | 4 | 2 | 2 | 2 | LLM consolidate differencet expressions into the same standard expresison |
| 14742428 | protein | {'S133sp', 'C219W', 'H391R', 'V69G', 'H156Y'} | {'C219W', 'H391R', 'V69G', 'H156Y', 'S133*'} | 5 | 5 | 4 | 1 | LLM use * instead of "sp" to represent a stop, which is more accurate |
| 15316799 | protein | {'F>S at codon 15'} | {'F15S'} | 1 | 1 | 0 | 1 | LLM format the unstructured text into a well-formmated variant |
| 16157158 | protein | {'C23S', 'C23S23'} | {'C23S'} | 2 | 1 | 1 | 1 | LLM consolidate differencet expressions into the same standard expresison |
| 16277682 | protein | {'methionine > valine substitution at codon404', 'M404V'} | {'M404V'} | 2 | 1 | 1 | 1 | LLM consolidate differencet expressions into the same standard expresison |

**Supplementary Table 5**. Disease benchmark and hallucination check using NCBI disease corpus. All of the LLM output pass manual examination for accuracy as well as hallucination. There is no hallucination and probably only 1 somehow “incorrect” disease extraction (PMID:1682919). The corresponding disease in the text has been highlighted in red font.

| **PMID** | **Disease_from_LLM** | **Max_Similarity_Score** | **Most_Similar_NCBI_Disease** | **Abstract (selected section)** | **Disease_Mentioned** | **Mnual Review** |
| --- | --- | --- | --- | --- | --- | --- |
| 10213492 | colorectal cancer | 0.666527 | colorectal adenomas | Inactivation of the adenomatous polyposis coli (APC) gene product initiates colorectal tumorigenesis. Patients with familial APC (FAP) carry germ-line mutations in the APC gene and develop multiple colorectal adenomas and subsequent carcinomas early in life. ... | FALSE | Pass |
| 10447259 | X-linked thrombocytopenia (XLT) | 0.619453 | XLT | Wiskott-Aldrich syndrome (WAS) is an X-linked recessive immunodeficiency characterized by thrombocytopenia, eczema, and recurrent infections, and caused by mutations in the WAS protein (WASP) gene. ... | FALSE | Pass |
| 1682919 | breast neoplasms | 0.592456 | B-cell neoplasms | ...; this highly conserved putative oncogene, which encodes a novel cyclin, has been linked to BCL1 and implicated also in subsets of breast and squamous cell neoplasms with 11q13 amplification. ... | FALSE | Pass (?); no hallucination |
| 1831007 | Huntington disease (HD) and Tourette syndrome (TS) | 0.696941 | Tourette syndrome | A 40-year-old man with childhood-onset Tourette syndrome (TS) developed Huntington disease (HD). We believe this to be the first reported case of childhood-onset TS with adult onset HD. Discovery of other cases with both disorders may provide clues to the pathophysiology of both conditions.. | FALSE | Pass; kept both disease at the same output |
| 1831007 | Tourette syndrome (TS) and Huntington disease (HD) | 0.705347 | Tourette syndrome | A 40-year-old man with childhood-onset Tourette syndrome (TS) developed Huntington disease (HD). We believe this to be the first reported case of childhood-onset TS with adult onset HD. Discovery of other cases with both disorders may provide clues to the pathophysiology of both conditions.. | FALSE | Pass |
| 2253937 | Becker muscular dystrophy (BMD) | 0.564284 | DMD | ... When used in conjunction with an existing primer set, these two multiplex reactions detect about 98% of deletions in patients with Duchenne or Becker muscular dystrophy (DMD, BMD). .. | FALSE | Pass |
| 2253937 | Duchenne muscular dystrophy (DMD) | 0.726816 | DMD | ... When used in conjunction with an existing primer set, these two multiplex reactions detect about 98% of deletions in patients with Duchenne or Becker muscular dystrophy (DMD, BMD)... | FALSE | Pass |
| 3169738 | Duchenne muscular dystrophy (DMD) and Becker muscular dystrophy (BMD) | 0.6159 | DMD | A panel of patients with Duchenne and Becker muscular dystrophy (DMD and BMD) has been screened with the cDNA probes Cf56a and Cf23a, which detect exons in the central part of the DMD gene... | FALSE | Pass |
| 3169738 | Becker muscular dystrophy (BMD) | 0.564284 | DMD | A panel of patients with Duchenne and Becker muscular dystrophy (DMD and BMD) has been screened with the cDNA probes Cf56a and Cf23a, which detect exons in the central part of the DMD gene. … | FALSE | Pass |
| 3169738 | Duchenne muscular dystrophy (DMD) | 0.726816 | DMD | A panel of patients with Duchenne and Becker muscular dystrophy (DMD and BMD) has been screened with the cDNA probes Cf56a and Cf23a, which detect exons in the central part of the DMD gene. ... | FALSE | Pass |
| 409732 | complement deficiency (specifically C7 deficiency) | 0.74344 | deficient in the seventh component of complement | In addition, the simultaneous occurrence of two hereditary complement deficiencies (C2 and C7) was discovered in one family of this remarkable kindred.. | FALSE | Pass |
| 409732 | complement deficiency (specifically C2 deficiency) | 0.674351 | deficient in the seventh component of complement | In addition, the simultaneous occurrence of two hereditary complement deficiencies (C2 and C7) was discovered in one family of this remarkable kindred.. | FALSE | Pass |
| 7573040 | spinocerebellar ataxia 3 (SCA3) | 0.735692 | SCA3 | The spinocerebellar ataxia 3 locus (SCA3) for type I autosomal dominant cerebellar ataxia (ADCA type I), a clinically and genetically heterogeneous group of neurodegenerative disorders, has been mapped to chromosome 14q32. 1 1. ... | FALSE | Pass |
| 7962532 | high HDL cholesterol levels | 0.429721 | cholesteryl ester transfer protein (CETP) deficiency | Genetic determinants of HDL cholesterol (HDL-C) levels in the general population are poorly understood. We previously described plasma cholesteryl ester transfer protein (CETP) deficiency due to an intron 14 G (+ 1) -to-A mutation (Int14 A) in several families with very high HDL-C levels in Japan. ... | FALSE | Pass |
| 8282802 | Classic (complete) lecithin cholesterol acyltransferase (LCAT) deficiency and Fish-eye disease | 0.768134 | cholesterol acyltransferase deficiency | Classic (complete) lecithin cholesterol acyltransferase (LCAT) deficiency and Fish-eye disease (partial LCAT deficiency) are genetic syndromes associated with markedly decreased plasma levels of high density lipoprotein (HDL) cholesterol but not with an increased risk of atherosclerotic cardiovascular disease. ... | FALSE | Pass |
| 8314592 | Norrie disease (pseudoglioma) | 0.769218 | Norrie disease | Positional cloning experiments have resulted recently in the isolation of a candidate gene for Norrie disease (pseudoglioma; NDP), a severe X-linked neurodevelopmental disorder. ... | FALSE | Pass |
| 8563759 | breast cancer | 0.535977 | ovarian cancers | The breast and ovarian cancer susceptibility gene, BRCA1, has been cloned and shown to encode a zinc-finger protein of unknown function. Mutations in BRCA1 account for at least 80% of families with both breast and ovarian cancer, as well as some non-familial sporadic ovarian cancers. ... | FALSE | Pass |
